# Epigenome-wide meta-analysis of PTSD across 10 military and civilian cohorts identifies novel methylation loci

**DOI:** 10.1101/585109

**Authors:** Alicia K Smith, Andrew Ratanatharathorn, Adam X Maihofer, Robert K Naviaux, Allison E Aiello, Ananda B Amstadter, Allison E Ashley-Koch, Dewleen G Baker, Jean C Beckham, Marco P Boks, Evelyn Bromet, Michelle Dennis, Sandro Galea, Melanie E Garrett, Elbert Geuze, Guia Guffanti, Michael A Hauser, Seyma Katrinli, Varun Kilaru, Ronald C. Kessler, Nathan A Kimbrel, Karestan C Koenen, Pei-Fen Kuan, Kefeng Li, Mark W Logue, Adriana Lori, Benjamin J Luft, Mark W Miller, Jane C Naviaux, Nicole R Nugent, Xuejun Qin, Kerry J Ressler, Victoria B Risbrough, Bart P F Rutten, Murray B Stein, Robert J. Ursano, Eric Vermetten, Christiaan H Vinkers, Lin Wang, Nagy A Youssef, Monica Uddin, Caroline M Nievergelt

## Abstract

Differences in susceptibility to posttraumatic stress disorder (PTSD) may be related to epigenetic differences between PTSD cases and trauma-exposed controls. Such epigenetic differences may provide insight into the biological processes underlying the disorder. Here we describe the results of the largest DNA methylation meta-analysis of PTSD to date with data from the Psychiatric Genomics Consortium (PGC) PTSD Epigenetics Workgroup. Ten cohorts, military and civilian, contributed blood-derived DNA methylation data (HumanMethylation450 BeadChip) from 1,896 PTSD cases (42%) and trauma-exposed controls (58%). Utilizing a common QC and analysis strategy, we identified ten CpG sites associated with PTSD (9.61E-07<p<4.72E-11) after adjustment for multiple comparisons (FDR<.05). Several CpGs were located in genes previously implicated in PTSD and other psychiatric disorders. The top four CpG sites fell within the aryl-hydrocarbon receptor repressor (*AHRR*) locus and were associated with lower DNA methylation in PTSD cases relative to controls. Interestingly, this association appeared to uncorrelated with smoking status and was most pronounced in non-smokers with PTSD. Additional evaluation of metabolomics data supported our findings and revealed that *AHRR* methylation associated with kynurenine levels, which were lower among subjects with PTSD relative to controls. Overall, this study supports epigenetic differences in those with PTSD and suggests a role for decreased kynurenine as a contributor to immune dysregulation in PTSD.

## Introduction

Posttraumatic stress disorder (PTSD) is characterized by re-experiencing, avoidance, and hyperarousal symptoms that can negatively impact mood and physiologic health ^1^. Although not everyone who experiences trauma goes on to develop PTSD, those who do often experience severe and disabling symptomatology ^2, 3^. Research suggests that both genetic and environmental factors contribute to risk for developing PTSD ^4, 5^.

Given the necessary, but not sufficient, role of environmental exposure (i.e., trauma) in the development of PTSD, it is critical to characterize the pathways underlying differential risk and resilience. Studies that contexualize the role of environmental influences provide additional insight into modifiable factors that may promote post-trauma resilience. For example, lack of social support at the time of trauma is associated with increased risk of developing PTSD in both military and civilian contexts ^6^. Similarly, studies that investigate how presumed environmental influences might affect biological pathways could provide insights into the genes whose regulation patterns differ in those with PTSD. A growing line of research aimed at elucidation of mechanisms by which environmental factors contribute to PTSD has focused on epigenetic modifications, which regulate gene function in response to environmental triggers. Epigenetic modifications, such as DNA methylation at cytosine-guanine dinucleotides (CpG sites), can induce changes in gene expression that are maintained through each round of cell division.

Multiple reviews have linked traumatic stress to differences in the proportion of methylated DNA (ß) at specific CpG sites ^7–10^. Indeed, a number of epigenome-wide association studies (EWAS) of individual cohorts have identified PTSD-associated CpG sites in genes and pathways related to neurotransmission and immune function ^11–15^. Similarly, studies using specific CpG sites that capture age acceleration demonstrate associations with PTSD and link differences in peripheral DNA methylation to memory formation and grey matter integrity ^16^. Although promising, the extant literature of EWAS studies on PTSD is limited by use of individual cohorts with small sample sizes and low statistical power, as well as the use of varying analytic pipelines that can make it challenging to synthesize findings. Consortia efforts can overcome these limitations by providing a shared analytic pipeline and increasing sample size and thus statistical power. The goal of this study is to capitalize on consortium strengths by conducting a meta-analysis of DNA methylation across 10 military and civilian cohorts participating in the Psychiatric Genomics Consortium (PGC) PTSD Epigenetics Workgroup.

## Materials and Methods

### Posttraumatic Stress Disorder Cohorts and Assessments

The participating cohorts, presented in Table 1, consisted of three civilian cohorts: the Detroit Neighborhood Health Study (DNHS), the Grady Trauma Project (GTP), and the World Trade Center 9/11 First Responders study (WTC); and seven military cohorts: the Army Study to Assess Risk and Resilience in Servicemembers (Army STARRS), the Marine Resiliency Study (MRS), the Injury and Traumatic Stress study (INTRuST), the Prospective Research in Stress Related Military Operations study (PRISMO), a European and African-American cohort from the Veterans Affairs’ Mental Illness Research, Education and Clinical Centers (VA-M), and a cohort from the National Center for PTSD (VA-NCPTSD). All subjects participating in these studies provided informed consent, and all studies were approved by respective institutional review boards. Current PTSD diagnosis was assessed by each individual cohort in accordance with the harmonization principles adopted by the PGC-PTSD Workgroup ^4^. Briefly, diagnosis of current PTSD was based on the diagnostic criteria defined by each cohort’s principal investigator (see cohort descriptions in Supplemental Material for complete details). All control subjects were trauma-exposed, and, if assessed in the respective cohort, control subjects that had a prior history of PTSD were excluded. Covariates were age, sex, genetic ancestry, and smoking status. A total of 1,896 subjects (42% cases) with DNA methylation from whole blood measured using the Illumina HumanMethylation450 BeadChip were selected for inclusion in this meta-analysis.

**Table 1.** Overview of the cohorts. Participating cohorts include: Detroit Neighborhood Health Study (DNHS), Grady Trauma Project (GTP), World Trade Center 9/11 Responders, SUNY (WTC), Army Study to Assess Risk and Resilience in Servicemembers (Army STARRS), Injury and Traumatic Stress (INTRuST), Marine Resiliency Study (MRS), Prospective Research in Stress-related Military Operations (PRISMO), Mid-Atlantic VA VISN 6 MIRECC (VA-M-AA & VA-M-EA), Boston VA/National Center for PTSD (VA-NCPTSD).

|  | **Study** | **N** | **PTSD (%)** | **Male (%)** | **Smokers (%)** | **White (%)** | **Black (%)** | **Hispanic (%)** | **Other or Unknown (%)** |
| --- | --- | --- | --- | --- | --- | --- | --- | --- | --- |
| Civilian | DNHS | 100 | 40% | 40% | 32% | 15% | 85% | 0% | 0% |
|  | GTP | 265 | 28% | 29% | 30% | 6% | 94% | 0% | 0% |
|  | WTC | 180 | 47% | 100% | 9% | 76% | 4% | 0% | 20% |
| Military | Army STARRS | 102 | 50% | 100% | 70% | 100% | 0% | 0% | 0% |
|  | MRS | 126 | 50% | 100% | 56% | 57% | 8% | 25% | 9% |
|  | INTRuST | 303 | 38% | 66% | 25% | 67% | 19% | 7% | 7% |
|  | PRISMO | 62 | 52% | 100% | 61% | 100% | 0% | 0% | 0% |
|  | VA-M-EA | 176 | 49% | 78% | 26% | 100% | 0% | 0% | 0% |
|  | VA-M-AA | 369 | 50% | 50% | 30% | 0% | 100% | 0% | 0% |
|  | VA-NCPTSD | 213 | 69% | 90% | 23% | 100% | 0% | 0% | 0% |
|  | Civilian Subtotal | 545 | 42% | 54% | 23% | 31% | 63% | 0% | 7% |
|  | Military Subtotal | 1351 | 42% | 56% | 34% | 61% | 32% | 4% | 3% |
|  | **Total** | **1896** | **42%** | **55%** | **31%** | **52%** | **41%** | **3%** | **3%** |

### Quality Control (QC) Procedures

Each study performed analyses at their site. To ensure consistent QC procedures across all participating cohorts, a set of common scripts were developed and implemented uniformly. This process is described by Ratanatharathorn and colleagues ^17^. Briefly, ß values, representative of the proportion of methylation at each probe, were caculated for all CpG sites in each sample. Samples with probe detection call rates <90% and those with an average intensity value of either <50% of the experiment-wide sample mean or <2,000 arbitrary units (AU) were excluded. Probes with detection p-values >0.001 or those based on less than three beads were set to missing as were probes that cross-hybridized between autosomes and sex chromosomes ^18^. CpG sites with missing data for >10% of samples within cohorts were excluded from analysis. Normalization of probe distribution was conducted using Beta Mixture Quantile Normalization (BMIQ) ^18^ after background correction. ComBat was used to account for sources of technical variation including batch and positional effects ^19^, while preserving variation attributable to study-specific outcomes and covariates that would be used in downstream analyses (e.g. case status or sex). Proportions of CD8, CD4, NK, B cells, monocytes, and granulocytes were estimated using each individual’s DNA methylation data based on the approach described by Houseman and colleagues and publicly available reference data (GSE36069) ^20, 21^.

### Statistical Analysis

Within each cohort, logit transformed ß values (M-values) ^22^ were modeled by linear regression as a function of PTSD, adjusting for sex (if applicable), age, CD8, CD4, NK, B cell, and monocyte cell proportions, and ancestry using principal components (PCs). For cohorts with available GWAS data (Army STARRS, GTP, INTRuST, MRS, VA-M, VA-NCPTSD), PCs 1-3 were included as covariates. For cohorts without GWAS data (DNHS, PRISMO, WTC), the method described by Barfield and colleagues was used to generate ancestry PCs directly from the methylation data, and PCs 2-4 were used to adjust for ancestry as recommended by the authors ^23^. Using the empirical Bayes method in the R package limma,^24^ moderated t-statistics were calculated for each CpG site, converted first into one-sided p-values then converted into z-scores to account for direction of effects. Meta-analysis across cohorts was performed by weighting each cohorts’ z-scores by its sample size relative to the total meta-analysis sample, summing weighted z-scores across cohorts from which two-sided p-values were calculated. P-values were adjusted for multiple-testing by controlling the False Discovery Rate (FDR) at 5 percent ^25^. MissMethyl was used to determine whether any of PTSD-associated CpGs were more likely to occur in Gene Ontology pathways than would be expected by chance alone; this approach has been optimized to account for the differences in the numbers of probes per gene and provide a more conservative multiple test correction for EWAS studies ^26^.

### Metabolite Analysis in the Marine Resiliency Study (MRS)

Targeted, broad-spectrum metabolomics was performed as previously described^27, 28^ with minor modifications. Lithium heparin plasma samples were collected and stored at −80°C until used for analyses in the MRS metabolomics study. Samples were analyzed from 116 participants exposed to military combat; 53 diagnosed with PTSD, and 63 controls without. Ninety microliters of plasma and 10 µl of added stable isotope internal standards were combined, extracted, and analyzed by hydrophilic interaction chromatography, electrospray ionization, and tandem mass spectrometry on a SCIEX 5500 QTRAP HILIC-ESI-MS/MS platform. From the 486 metabolites measured, we extracted and analyzed data for kynurenine and kynurenic acid to test our hypothesis that *AHRR* methylation associates with trypophan breakdown products. Cotinine, a metabolite of nicotine with a plasma half-life of about 18 hours, was measured to provide independent information about tobacco exposure. A cotinine area under the curve (AUC) of 1 × 10^6^ was used as the upper limit for a non-smoker or second-hand tobacco exposure. Analyte AUC data was log-transformed, and analyte z-scores were used for further statistical analysis. To determine if trytophan metabolite levels differed between PTSD cases and controls, we performed linear regressions of metabolites on PTSD status, including GWAS PCs 1-3 and cell type proportions as covariates. To determine if tryptophan metabolite levels were associated with *AHRR* methylation, we performed linear regressions of metabolites on each *AHRR* CpG site, including PCs and cell type proportions as covariates. Correlation between cotinine and kynurenine was measured using Pearson’s correlation coefficient.

## Results

### Participating cohorts

Sample characteristics for the 10 cohorts that have contributed data are listed in Table 1 (N=1,896). All participants were exposed to trauma, and 42% had a current diagnosis of PTSD. There were no significant age differences in PTSD cases and trauma-exposed controls. However, the demographic characteristics for each cohort varied substantially (Table S1), with cases more likely to be male and current smokers (p < .05) across the majority of cohorts and in the overall cohort.

### PTSD-associated CpG sites from Meta-Analysis

In our primary analysis we found 10 CpG sites significantly associated with current PTSD (9.61E-07<p<4.72E-11; Figure 1; Table 2) that remained significant after controlling for multiple comparisons (FDR<.05). The aryl hydrocarbon receptor repressor (*AHRR*) contains the top 4 PTSD-associated CpGs, with lower methylation in PTSD cases relative to controls (Figure S1). Though methylation of these CpGs are moderately correlated, they are located over ~22kb from each other. Some of the other PTSD-associated CpG sites (Table 2) were located in genes involved in other epigenetic processes, including microRNA and noncoding RNAs (e.g. *MIR3170, AC011899.9* and *LINC00599*).

**Table 2.** CpG sites significantly associated with current PTSD (FDR<0.05).

| **CpG** | **Location** | **Gene** | **Features** | **Z** | **p-value** | **FDR** |
| --- | --- | --- | --- | --- | --- | --- |
| cg05575921 | chr5:373378 | *AHRR* | Body | 6.58 | 4.72E-11 | 2.15E-05 |
| cg21161138 | chr5:399360 | *AHRR* | Body | 6.24 | 4.39E-10 | 1.00E-04 |
| cg25648203 | chr5:395444 | *AHRR* | Body | 5.72 | 1.07E-08 | 1.63E-03 |
| cg26703534 | chr5:377358 | *AHRR* | Body | 5.59 | 2.25E-08 | 2.57E-03 |
| cg25415650 | chr13:26795862 | *RNF6* | 5'UTR | -5.38 | 7.38E-08 | 6.73E-03 |
| cg17284326 | chr13:98749760 | *MIR3170* | IGR | 5.35 | 8.93E-08 | 6.79E-03 |
| cg07339236 | chr20:50312490 | *ATP9A* | Body | 5.17 | 2.39E-07 | 1.56E-02 |
| cg26801037 | chr7:157294357 | *AC011899.9* | IGR | 5.12 | 3.08E-07 | 1.76E-02 |
| cg14405344 | chr9:84609982 | *FLJ46321* | Body | 4.94 | 7.86E-07 | 3.99E-02 |
| cg18217048 | chr8:9742024 | *LINC00599* | IGR | 4.90 | 9.61E-07 | 4.39E-02 |

**Figure 1.**
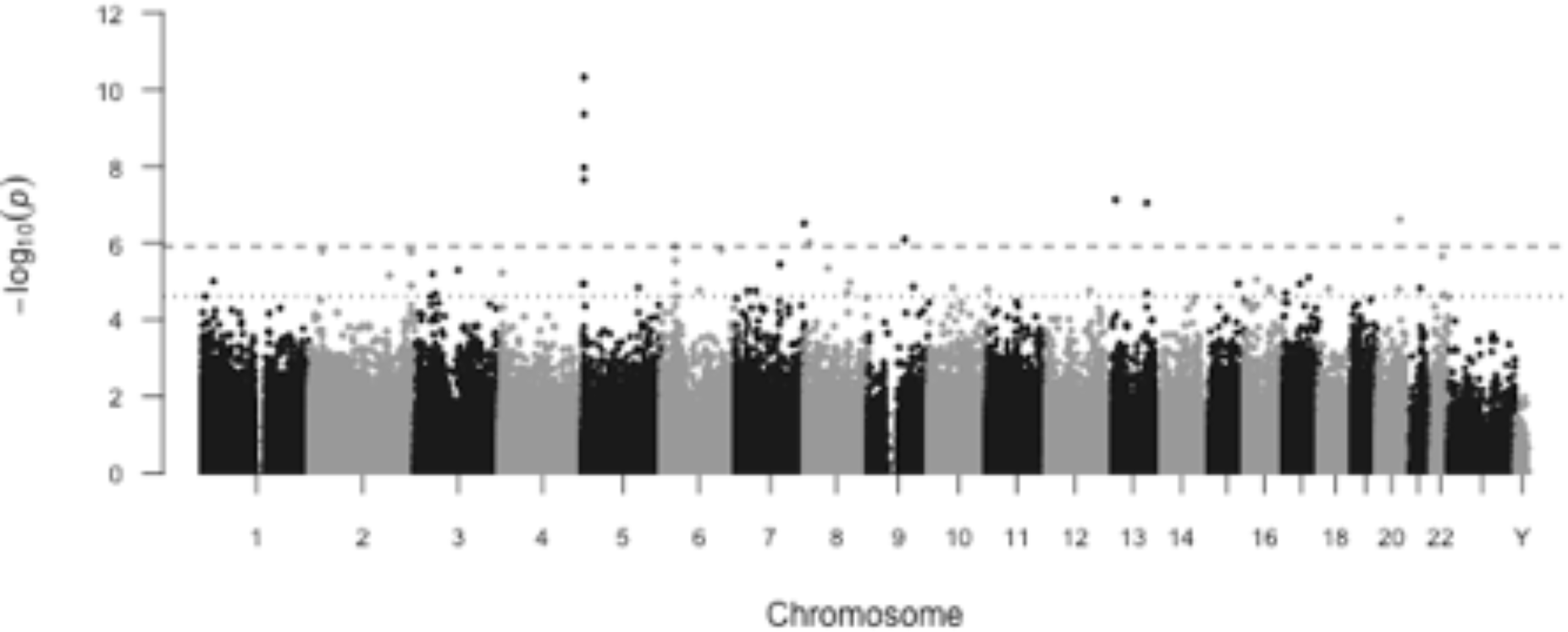
PTSD associates with DNA methylation differences across the genome. A Manhattan plot of the association of current PTSD with DNA methylation proportion. The x-axis is the location of each site across the genome. The y-axis is the –log10 of the p-value for the association with PTSD. The top dashed line indicates statistical significance at FDR < 0.05, and the lighter dashed line below indicates statistical significance at FDR < 0.2.

### Sensitivity Analysis with Smoking Status

Because lower methylation of *AHRR* CpG sites has been associated with smoking ^29–31^, we controlled for smoking status in our sensitivity analyses of the 10 significant CpGs (Figure S2). The association between *AHRR* methylation and PTSD was substantially attenuated for all four CpGs; the attenuation effect was weaker for the six CpGs in other genes. However, since the higher rates of smoking among PTSD cases introduced the possibility of collinearity, we evaluated the association between methylation of the top 10 CpGs and PTSD separately in smokers and non-smokers (Figure 2). While some CpGs were comparably associated in both smokers and non-smokers (e.g. cg25415650 in *RNF6* or cg17284326 in *MIR3170*), the associations between *AHRR* CpGs and PTSD were most prominent in non-smokers.

**Figure 2.**
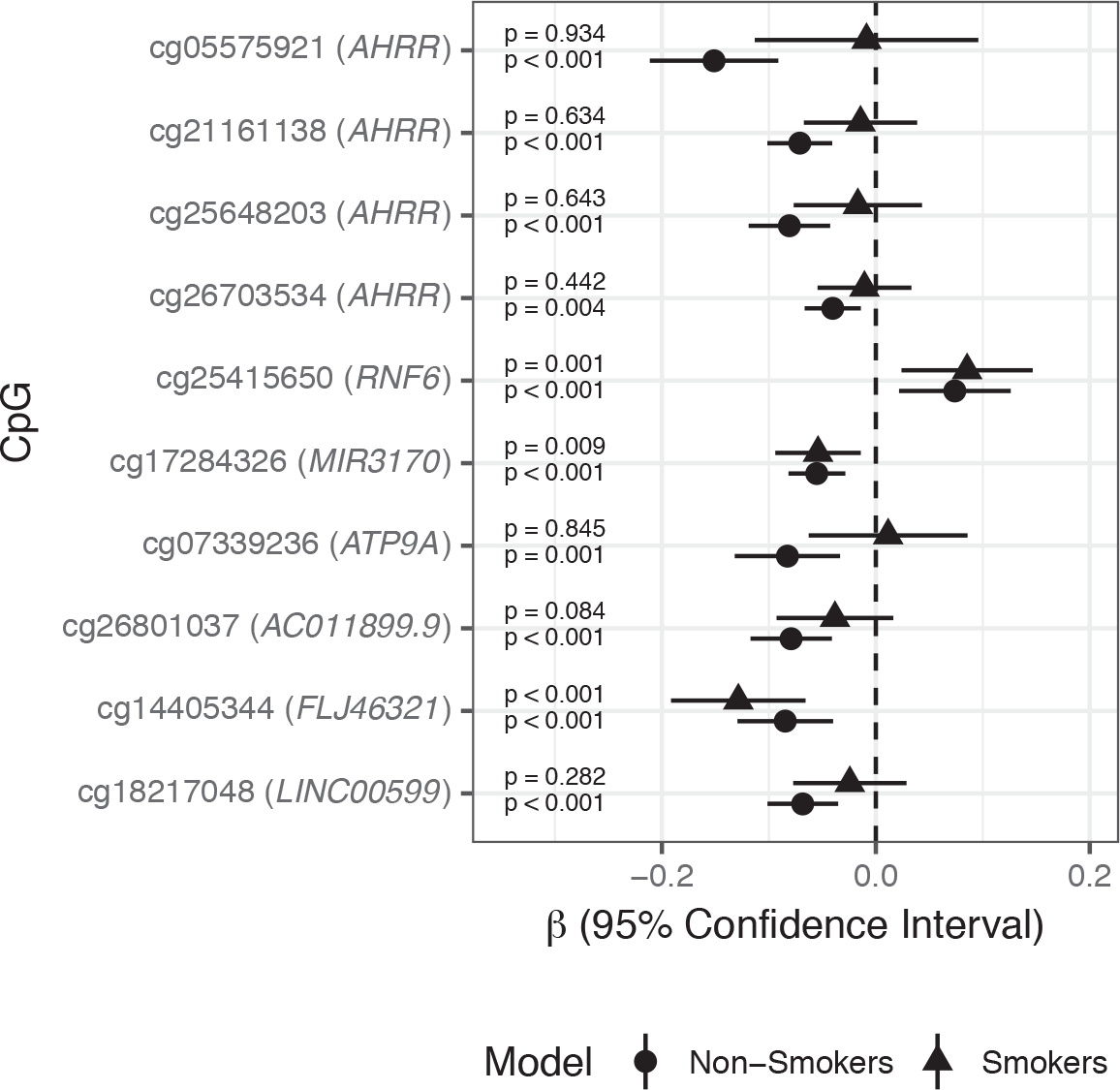
Associations between PTSD-associated CpGs (FDR<.05) stratified by smoking status. The x-axis is the effect size for each association and includes the 95% confidence interval.

We further delineated the association between PTSD and smoking-associated CpG sites with the hypothesis that un-reported smoking among those that identify as non-smokers would result in the appearance of an association between smoking-associated CpG sites and PTSD, leveraging information from a large meta-analysis on smoking conducted by the CHARGE consortium ^31^. This evaluation included 21 CpGs representing the most significantly associated CHARGE smoking sites and CpGs in *AHRR*. If the association between *AHRR* and PTSD was due to unreported smoking in PTSD controls, we would expect to see a similar effect across all smoking-CpGs. However, after adjusting for the number of tests performed, 3 *AHRR* CpG sites remained associated with PTSD in the non-smokers only, and no smoking CpG showed a significant association with PTSD in the controls or cases. These findings are consistent with the hypothesis that the association between PTSD and DNA methylation of *AHRR* is independent of smoking status (Figure S3).

### Tryptophan Catabolism in PTSD

We next evaluated other biological factors that could contribute to aryl-hydrocarbon receptor (AHR) expression. AHR is activated in many immune cell types, including T cells, B cells, and NK cells, by indolamine-mediated tryptophan catabolism ^32^. Inflammation can shift tryptophan metabolism away from serotonin synthesis towards kynurenine synthesis ^33^. To evaluate the association of tryptophan catabolism in PTSD and its relationship to *AHRR* methylation, we leveraged data from the 116 subjects from MRS that had both DNA methylation and tryptophan metabolite data. In this group, we first note that kynurenine levels were lower in subjects with PTSD relative to controls (p=0.048; Figure S4). Consistent with this observation, lower methylation of *AHRR* CpGs associated with lower kynurenine (cg21161138; p=0.039) and lower kynurenic acid (cg05575921; p=0.004).

Further, metabolome data in the MRS confirmed that self-reported smoking status is consistent with empiric cotinine levels; 87% of self-reported non-smokers and light smokers had cotinine levels consistent with no, second hand or light exposure (i.e. cotinine AUC < 1 × 10^6^) while 91% of regular or heavy smokers had cotinine levels consistent with their endorsement (i.e. cotinine AUC ≥ 1 × 10^7^). In addition, these data also provide insight into why controlling for smoking may attenuate the association between DNA methylation and PTSD. We noted an inverse relationship between cotinine and kynurenine (p=0.0004) that was consistent among both smokers and non-smokers (Figure S5).

### Gene Ontology enrichment analysis

Evaluation of the 50 CpG sites in 42 genes or non-coding RNAs associated with PTSD (FDR<.2; Additional File 1) revealed that several were located in genes previously implicated in psychiatric disorders or pharmacologic treatment response (e.g. *AGBL1, ATP9A, CUX1, FLJ46321* aka *SPATA31D1, GRIN3A, GOT2, HOXA3, NEUROD2*, and *SYNJ1*). However, gene ontology analysis did not support enrichment of PTSD-associated CpGs in any biological processes after correction for multiple comparisons (Table S2).

## Discussion

An individual’s risk of developing PTSD depends on both the nature of the trauma and the physiological response to that trauma ^34^. Not all individuals exposed to trauma develop PTSD, and a better understanding of the modifiable biological factors underlying risk and resilience will inform the development of new prevention and treatment strategies. In this study we identified CpG sites associated with PTSD, some of which occur in genes implicated in previous studies of PTSD (e.g. *MBL2*, *NCF4*, and *TMEM49*) ^35, 36^ and pharmacologic response to medications used in PTSD treatment (e.g. *CUX1*, *FARP1*, *GRIN3A*, and *MIR3170*) ^37–39^. We identified PTSD-associated CpGs in genes implicated in other neuropsychiatric disorders (e.g. *AGBL1*, *ATP9A*, *FLJ46321*, *GOT2*, *HOXA3*, *MAP7*, and *NEUROD2*) ^40–45^, which may provide insight into comorbidities commonly reported in PTSD. We also identified CpGs in genes that are involved in other epigenetic mechanisms (e.g. *AC011899.9*, *C20orf199*, *LINC00599*, *MIR548A3*, *RP11-290F5.2*, and *SNHG5*). Numerous studies in the blood of subjects with PTSD report differences in microRNA (miRNA) and noncoding RNAs (ncRNA) ^46–49^.

However, the most interesting finding from this study was that, on average, PTSD cases, relative to controls, had lower methylation at several CpG sites in the *AHRR* gene when compared to trauma-exposed controls. Methylation of *AHRR* CpGs has been strongly linked to smoking ^29–31^. As substantially more of the PTSD cases reported smoking compared to controls, this suggested that we should control for smoking in the EWAS. A comparable approach was taken by Marzi and colleagues ^50^. In their study of DNA methylation in relation to victimization stress in adolescents, they noted experiment-wide significant associations in 3 of the 4 *AHRR* CpGs associated with PTSD. Similar to our findings, the association of *AHRR* and PTSD was no longer significant after statistically controlling for smoking. Marzi and colleagues concluded that there are no robust changes in DNA methylation in victimized young people. Our study went beyond this approach to conduct a stratified analysis that revealed the most prominent PTSD-associated difference in *AHRR* methylation was evident in the non-smokers. To evaluate the possibility of confounding due to un-reported smoking among those that identify as non-smokers, we tested top smoking-related CpGs, including *AHRR*, for association with PTSD, and identified no evidence of association between smoking-related CpG sites and PTSD, supporting our conclusion that the association between *AHRR* CpG sites and PTSD was independent of smoking.

Over the last decade, there have been a wealth of studies describing the role of the aryl hydrocarbon receptor in the immune system, with specific roles in T cells, B cells, monocytes and dendritic cells (DC) ^32, 51–53^. The aryl hydrocarbon receptor (AhR) pathway works in a regulatory capacity. Briefly, when a ligand binds the AhR, it translocates to the nucleaus where it drives expression of its target genes, including the aryl hydrocarbon receptor repressor (AHRR), which begins a feedback loop in which it can competitively bind the AhR. The AhR pathway can either limit or stimulate an inflammatory response. One mechanism by which this occurs is by promoting differentiation of T cells into T regulatory cells (Tregs) or T helper 17 (Th17), though this appears to be done in a ligand-specific manner. For example, endogenous ligands, such as kynurenine or dietary indoles^54^, tend to promote differentiation into Tregs, which reduce the immune response in a self-limiting manner, resulting in reduced inflammation and less Treg generation. In contrast, exogenous ligands, such as dioxin or polyaromatic hydrocarbons in cigarette smoke, tend to promote differentiation into Th17, which promotes a heightened inflammatory response ^51^. However, most studies of the AhR pathway stimulate it with a single, typically exogenous, ligand, making it difficult to interpret how this biology may be relevant to specific conditions that may involve stimulation by both ligand types. Recent studies also suggest that different groups of genes are expressed based on the different ligand types. For example, kynurenine binding to AhR stimulates the expression of anti-inflammatory genes such as *TGF-β* and *IDO1*, while dioxin (2,3,7,8-tetrachlorodibenzo-ρ-dioxin; TCDD) binding to the AHR preferentially activates the drug metabolizing enzymes, such as *CYP1A1* and *CYP1B1^55^*, and pro-inflammatory cytokines, such as IL1-*β*, IL2, IL6, and TNF-α^56^. A recent study shows that, when stimulated by TCDD, AHRR selectively inhibits only subsets of genes that are activated by AHR, further suggesting that the epigenetic activation of *AHRR* may have selective actions on anti-inflammatory and pro-inflammatory actions in immune cells ^57^.

In a subset of subjects with tryptophan metabolite data, we reported that kynurenine levels were significantly lower in subjects with PTSD relative to trauma-exposed controls and that lower methylation of *AHRR* CpGs associated with lower kynurenine and its metabolites. This pattern is similar to what is observed following chronic exposure to nicotine and possibly other AHR-stimulating ligands ^58^. This chronic exposure scenario is reflected in our cohort among both controls and PTSD cases by the finding that higher levels of cotinine were associated with decreased levels of kynurenine. Independent of its source, lower levels of kynurenine would likely result in a counter-regulatory increase in Th17 pro-inflammatory activity ^59^, and may help to account for the frequent observation of heightened inflammatory activity observed in subjects with PTSD ^60^.

While smokers are more common among our PTSD group, approximately half of the cases in our cohorts are non-smokers, who exhibited the most prominent associations between *AHRR* CpGs and PTSD. Methylation of *AHRR* CpGs decreases only so much even among heavy smokers suggesting that there may be limited variability in these subjects and a limited degree to which methylation at this site may be decreased. Though the vast majority of the *AHRR* methylation literature is focused on characterizing its variation in the context of smoking, the types of polycyclic aromatic hydrocarbons that stimulate AHR are present from multiple common sources including burning wood or charcoal, auto emissions, industrial exhaust, and urban dust ^61^. Similarly, participants in the WTC and military cohorts are likely to have substantial levels of occupational exposure that could result in AHR activation ^62–64^. In this context, it is reasonable to hypothesize that smoking is just one of many potential environmental exposures that could promote inflammation following AHR stimulation (Figure 3).

**Figure 3.**
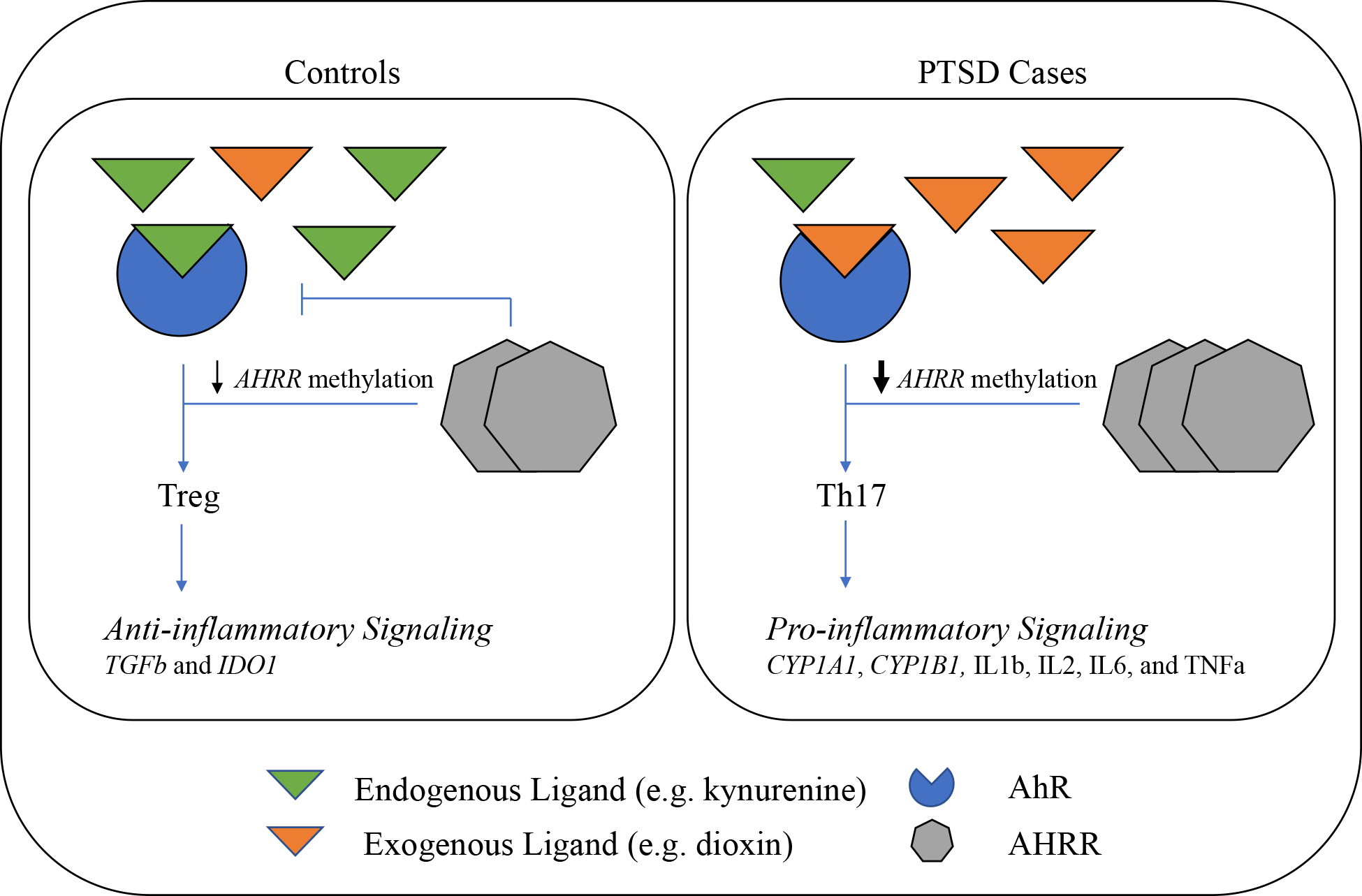
Conceptual Model for *AHRR*-mediated immune dysregulation in PTSD. Our data suggests that PTSD cases have more exposure to exogenous AhR ligands through behavioral (i.e. smoking) and occupational (i.e. airborne particulates) sources, resulting in persistently lower *AHRR* methylation, higher *AHRR* expression, and AhH-mediated pro-inflammatory signaling. Activation of AhR signaling in Controls is more likely to result from endogenous ligand binding (i.e. kynurenine or dietary products), which promotes a transient decrease in *AHRR* methylation, anti-inflammatory signaling, and negative regulation of AhR.

Both epidemiologic and immunologic studies report immune dysregulation in those with PTSD compared to controls. For example, autoimmune and inflammatory disorders, such as rheumatoid arthritis, have been linked to PTSD ^65–67^. Studies of the immune system generally support higher levels of pro-inflammatory cytokines in PTSD cases relative to controls ^60^. A study investigating Tregs reported a lower number of Tregs in the blood of PTSD patients ^68^ while another reported a difference in the composition of Treg populations that suggested higher susceptibility for autoimmunity ^69^. Another study reported lower proportions of Tregs and higher proportions of Th17 cells in PTSD cases before linking Th17 counts to higher clinical symptom scores ^48^. Finally, a randomized control trial of subjects undergoing narrative exposure psychotherapy noted higher levels of Tregs and lower PTSD symptoms following treatment ^70^.

This study has a number of limitations that should be considered. First, we used existing data generated on the HumanMethylation450 array, which captures only a fraction of the CpG sites in the genome. Though sequencing methods would capture a larger proportion of the epigenome that may include regions important for PTSD, focusing on this array allowed us to evaluate a larger and more diverse cohort of subjects and enabled processing and analysis of these diverse samples with a common analytic pipeline. Second, our study examined whole blood-derived DNA. Though this approach likely captures part of the PTSD sequelae and may be informative for future biomarker studies, it is unlikely to reflect DNA methylation in brain regions most relevant for PTSD. As studies of tissues from PTSD Brain Banks are conducted, it will be important to look for parallels between these different tissues. Third, our secondary analysis of DNA methylation and kynurenine was limited to the only cohort with cotinine and metabolomics data available on the subjects included in the methylation study, which limited sample size, and power for this portion of the study, and potentially generalizability to other cohorts. However, we believe that the ability to verify self-reported smoking status with a biological measure and to evaluate our kynurenine hypothesis was a considerable strength of the study. We hope these results will prompt other investigators to replicate these findings in their own cohorts. Fourth, we were unable to directly measure or impute the proportion of different types of T cells in this study. Though the degree to which the immune system is involved is speculative without functional studies, involvement of AHRR is supported by metabolomic data. Fifth, there is phenotypic heterogenetity among the cohorts contributing to this meta-analysis. Some cohorts assess individuals that were recently exposed to trauma while others include indivduals with chronic PTSD. Sixth, few of the cohorts included in this meta-analysis have detailed physical or psychiatric information on subjects prior to trauma exposure, making it difficult to evaluate the role of comorbidities. Thus, it is possible that some of the epigenetic differences observed in this study may have been in place prior to PTSD development. Finally, this meta-analysis uses only cross-sectional data. In order to establish whether these PTSD-associated differences are a cause or consequence of the disorder or both, longitudinal and experimental studies will be required.

Taken together, the results of this study implicates the immune system in PTSD and suggest that epigenetic mechanisms may play a role in that process. A substantial fraction of those diagnosed with PTSD do not respond to pharmacologic or psychological interventions, and clinical and preclinical studies have begun to evaluate strategies to limit inflammation as a first line treatment ^71, 72^. Future studies should evaluate the role of ligand-specific AHR activation in the development and progression of PTSD.

## Acknowledgements

This work was supported by the U.S. Army Medical Research and Materiel Command and the National Institute of Mental Health (NIMH; R01MH108826; R01MH106595) as well as the Biomedical and Laboratory Research and Development (#I01BX002577). We appreciate the technical support of all of the staff, volunteers and participants from the Grady Trauma Project, supported by the National Institutes of Mental Health (MH096764 and MH071537). DNHS, which is grateful to all of the participants and staff for their contributions, was funded by NIH Awards R01DA022720, R01DA022720-S1, and RC1MH088283. The Marine Corps, Navy Bureau of Medicine and Surgery (BUMED) and VA Health Research and Development (HSR&D) provided funding for MRS data collection and analysis and NIH R01MH093500 funded the GWAS assays and analysis. Acknowledged are Mark A. Geyer (UCSD), Daniel T. O’Connor (UCSD), all MRS investigators, as well as the MRS administrative core and data collection staff listed in the Methods article (Baker et al, Prev Chronic Dis. 2012;9(10):E97). The authors also thank the Marine and Navy Corpsmen volunteers for military service and participation in MRS. Support for metabolomics in the Naviaux lab at UCSD was provided in part by philanthropic gifts from the UCSD Christini Fund, the Lennox Foundation, the Wright Family Foundation, and the Jane Botsford Johnson Foundation. Data collection of PRISMO was funded by the Dutch Ministry of Defence, and DNA methylation analyses were funded by the VENI Award fellowship from the Netherlands Organisation for Scientific Research (NWO, grant number 916.11.086). The VA Boston-National Center for PTSD Study research was supported in part by National Institute of Mental Health Award RO1MH079806, Department of Veterans Affairs, Clinical Science Research & Development Program Award 5I01CX000431-02, Department of Veterans Affairs, Biomedical Laboratory Research & Development Program Award 1I01BX002150-01. This research is the result of work supported with resources and the use of facilities at the Pharmacogenomics Analysis Laboratory, Research and Development Service, Central Arkansas Veterans Healthcare System, Little Rock, Arkansas. This work was also supported by a Career Development Award to E. J. Wolf from the Department of Veterans Affairs, Clinical Sciences Research, and Development Program. Dr. Kimbrel was supported by a Career Development Award (#IK2CX000525) from the Clinical Science Research and Development (CSR&D). Dr. Beckham was supported by a Research Career Scientist Award (#11S-RCS-009) from the CSR&D Service of VA ORD. This research was also supported, in part, by a Merit Award (#I01BX002577) from the Biomedical Laboratory Research and Development (BLR&D) Service of VA ORD. The VA Mid-Atlantic Mental Illness Research, Education, and Clinical Center Workgroup includes John A. Fairbank, Mira Brancu, Patrick S. Calhoun, Eric A. Dedert, Eric B. Elbogen, Kimberly T. Green, Robin A. Hurley, Angela C. Kirby, Jason D. Kilts, Christine E. Marx, Gregory McCarthy, Scott D. McDonald, Marinell Miller-Mumford, Scott D. Moore, Rajendra A. Morey, Jennifer C. Naylor, Treven C. Pickett, Jared Rowland, Jennifer J. Runnals, Cindy Swinkels, Steven T. Szabo, Katherine H. Taber, Larry A. Tupler, Elizabeth E. Van Voorhees, H. Ryan Wagner, Richard D. Weiner, and Ruth Yoash-Gantz. Army STARRS was sponsored by the Department of the Army and funded under cooperative agreement number U01MH087981 (2009-2015) with the National Institutes of Health, National Institute of Mental Health (NIH/NIMH). WTC study was sponsored by CDC/NIOSH award U01 OH010416-01. The PTSD and TBI INjury and TRaUmatic STress Clinical Consortium (INTRuST) was funded by a grant from the United States Department of Defense: W81XWH08-2-0159. Members of the INTRuST Consortium Biorepository Working Group who contributed to this work include: Gerald A. Grant MD, Christine E. Marx MD, Mark S. George MD, Thomas W. McAllister MD, Norberto Andaluz MD, Lori Shutter MD, Raul Comibra MD, Ross D. Zafonte DO, Sonia Jain PhD, Xue-Jun Qin, and Michael Hauser PhD.

The views expressed in this article are those of the authors and do not necessarily reflect the position or policy of the VA, NIH, or the United States government.

