## Supplementary Materials for "Epigenome-wide meta-analysis of PTSD across 10 military and civilian cohorts identifies novel methylation loci"

**Supplemental Material**

Cohort Descriptions

*Detroit Neighborhood Health Study (DNHS)* - As detailed in a previous study [1], participants (n=1,547 at the baseline wave) were assessed for PTSD symptoms using the PTSD checklist (PCL-C), a 17-item self-report measure of Diagnostic and Statistical Manual of Mental Disorders (DSM-IV) symptoms, and additional questions about duration, timing, and impairment or disability due to the symptoms [2]. Participants were initially asked to identify Potentially Traumatic Events that they experienced in the past from a list of 19 events. PTSD symptoms were then assessed by referencing two traumatic events that the respondent may have experienced: one that the participant regarded as the worst and one randomly selected event from the remaining PTEs a respondent may have experienced. Respondents were considered affected by lifetime PTSD if all six DSM-IV criteria were met in reference to either the worst or the random event. The DNHS was approved by the institutional review board at the University of Michigan and University of North Carolina at Chapel Hill.

*Grady Trauma Project (GTP)* –GTP research participants were approached in the waiting rooms of the primary care clinic or obstetrical-gynecological clinic of a large, urban, public hospital in Atlanta, GA while either waiting for their medical appointments or while waiting with others who were scheduled for medical appointments [3]. Subjects willing to participate provided written informed consent and participated in a verbal interview and blood draw. Current and lifetime PTSD diagnosis was assessed by clinical psychologists using the Clinician-Administered PTSD Scale for DSM IV (CAPS-4) [4] or the Mini International Neuropsychiatric Interview DSM IV (MINI), an instrument designed to assess major Axis 1 disorders with high validity and reliability [5, 6]. For this study, cases were specified as having current PTSD, and controls had no current or lifetime history of the disorder. Demographic variables including age, sex and race were assessed through self-report. The Institutional Review Boards of Emory University School of Medicine and the Research Oversight Committee of Grady Memorial Hospital approved this study.

*World Trade Center Responders (WTC)* - As described by Bromet and colleagues [7, 8], responders enrolled in the Stony Brook University/Long Island WTC Health Program were administered the SCID PTSD module modified to assess PTSD in relation to WTC exposures. The sample providing blood samples for the epigenetics assays was restricted to men (the vast majority of the responders) and oversampled for posttraumatic stress disorder (PTSD) [9]. The Committees on Research Involving Human Subjects at Stony Brook University approved the study.

*Army Study to Assess Risk and Resilience in Servicemembers (Army STARRS):* Details of the diagnostic procedures are described in detail in prior STARRS publication [10]. Briefly, participants completed a computerized version of the Composite International Diagnostic Interview screening scales (CIDI-SC) and a screening version of the PTSD Checklist (PCL) for DSM-IV. Trauma exposure was assessed from answers pertaining to childhood, adulthood civilian, and military traumatic events. PTSD diagnosis was assigned using multiple imputation methods that relied on PCL and CIDI-SC data; our clinical reappraisal study found satisfactory concordance with independent clinical diagnoses based on blinded Structured Clinical Interviews for DSM-IV (AUC = 0.70–0.79; κ =0.4–0.6). Healthy controls were determined not to have PTSD using the same instruments and approach.

*Injury and Traumatic Stress (INTRuST)*: Patients and healthy controls were recruited from INTRuST clinical trials (with DNA collected prior to treatment) or specifically to contribute to the INTRuST biorepository. Patients were diagnosed with PTSD using study-specific measures which could have included either the CAPS-IV or CAPS-5, the MINI, or self-report using the PCL. Healthy controls were determined to be free of major psychiatric diagnoses, including PTSD, using the MINI or a comparable clinical interview. The Human Research Protection Program (HRPP) at UCSD approved this study, as did all the IRBs at participating sites.

*Marine Resiliency Study (MRS)* - In the MRS [11, 12], PTSD was diagnosed up to 3 times, once before deployment and 3 and/or 6 month post deployment. Post-traumatic stress (PTS) symptoms were assessed using a structured diagnostic interview, the Clinician Administered PTSD Scale (CAPS), and PTSD diagnosis followed the DSM-IV criteria for partial and full PTSD and a minimum CAPS score of 40. None of the participants in the EWAS and metabolome study had PTSD at pre-deployment. Samples (DNA from whole-blood and Li-heparin plasma for metabolome) of PTSD cases were selected from the 3- or 6-months post-deployment visits, choosing the visit with the highes CAPS score. Combat-exposed controls with low to no PTSD-symptoms were selected from matching post-deployment visits. The study was approved by the University of California – San Diego Institutional Review Board.

*Prospective Research in Stress-related Military Operations (PRISMO)* - All subjects in the DD were male participants in PRISMO, a large prospective study of 1,032 well-characterized Dutch military soldiers scheduled for a deployment of at least four months to Afghanistan with longitudinal follow-up. Baseline measures were recorded at one month before deployment. Follow-up was performed at one month and six months post-deployment, and data from the baseline and six-month follow-up were used for this analysis. A subset (total n=93) of three similarly sized subgroups of PRISMO study participants were pre-selected based on the level of traumatic stress exposure and the presence of PTSD symptoms: i) a subgroup showing high combat-trauma exposure (7.3 ± 2.9) and high levels of post-deployment PTSD symptoms (45.3 ± 8.6); ii) a subgroup showing high combat-trauma exposure (8.6 ± 2.3) and a low severity of PTSD symptoms (26.0 ± 3.7); and iii) a subgroup showing low combat-trauma exposure (0.4 ± 0.5) and low levels of post-deployment PTSD symptoms (25.1 ± 3.7).

Blood samples were collected six months after deployment. The blood cell-type composition was investigated using flow cytometry, implemented in the clinical laboratory of Utrecht University Medical Center, as previously reported [13]. The presence and severity of symptoms of PTSD over the previous four weeks were assessed with the 22-item Self-Report Inventory for PTSD (SRIP), which has good reliability and validity. Differences in PTSD symptoms between time points were log-transformed to improve the distribution. Exposure to traumatic stress during deployment was assessed with a 19-item deployment experiences checklist, as previously reported [14].

*Mid-Atlantic Mental Illness Research Education and Clinical Center PTSD Study (VA-M-AA & VA-M-EA)* - As described previously [15], PTSD was diagnosed using the Structured Clinical Interview for DSM-IV Disorders (SCID) administered by trained interviewers. In accordance with the DSM-IV, PTSD consists of three symptom clusters. These include re-experiencing symptoms (B symptoms), avoidance and numbing symptoms (C symptoms) and hyperarousal symptoms (C symptoms). Total PTSD symptoms and symptom clusters (B, C, or D) were measured using the Davidson Trauma Scale for all veterans including individuals with current PTSD diagnosis and controls. The research was reviewed and approved by the Institutional Review Boards at the Salisbury VA, Hampton VA, Durham VA and Duke University Medical Centers.

*Boston VA National Center for PTSD (VA-NCPTSD)* – As described by Logue and colleagues [16], VA study participants were administered the CAPS, a 30-item structured diagnostic interview that assesses the frequency and severity of the 17 DSM-IV PTSD symptoms, 5 associated features and functional impairment, to assess current and lifetime PTSD symptoms. The Institutional Review Boards at two VA health care facilities approved the study.

**Table S1. Detailed description of phenotypes by cohort.**

| | Cases | | | Control | | | Total | | |
|---|---|---|---|---|---|---|---|---|---|
| **Age** | **Mean (SD)** | | | **Mean (SD)** | | | **Mean** | | |
| DNHS | 50 (13.23) | | | 56 (14.11) | | | 53.6 | | |
| GTP | 38.88 (11.5) | | | 43.14 (12.52) | | | 41.95 | | |
| MRS | 22.16 (2.3) | | | 22.25 (3.65) | | | 22.2 | | |
| PRISMO | 26.75 (9.57) | | | 27.47 (9.01) | | | 27.1 | | |
| STARRS | 23.94 (4.93) | | | 23.63 (3.48) | | | 23.79 | | |
| TRUST | 38.11 (10.98) | | | 31.59 (11.43) | | | 34.09 | | |
| VA-M-AA | 38.16 (9.11) | | | 38.56 (9.62) | | | 38.36 | | |
| VA-M-EA | 34.53 (9.5) | | | 35.21 (10.3) | | | 34.87 | | |
| VA-NCPTSD | 31.86 (8.45) | | | 33.4 (10.06) | | | 32.33 | | |
| WTC | 48.39 (7.23) | | | 50.88 (8.93) | | | 49.72 | | |
| | **Cases** | | | **Control** | | | **Total** | | |
| **Gender** | **Male** | **Female** | **Missing** | **Male** | **Female** | **Missing** | **Male** | **Female** | **Missing** |
| DNHS | 13 | 27 | 0 | 27 | 33 | 0 | 40 | 60 | 0 |
| GTP | 19 | 55 | 0 | 59 | 132 | 0 | 78 | 187 | 0 |
| MRS | 63 | 0 | 0 | 63 | 0 | 0 | 126 | 0 | 0 |
| PRISMO | 32 | 0 | 0 | 30 | 0 | 0 | 62 | 0 | 0 |
| STARRS | 51 | 0 | 0 | 51 | 0 | 0 | 102 | 0 | 0 |
| TRUST | 102 | 14 | 0 | 99 | 88 | 0 | 201 | 102 | 0 |
| VA-M-AA | 92 | 91 | 0 | 93 | 93 | 0 | 185 | 184 | 0 |
| VA-M-EA | 67 | 20 | 0 | 71 | 18 | 0 | 138 | 38 | 0 |
| VA-NCPTSD | 128 | 20 | 0 | 63 | 2 | 0 | 191 | 22 | 0 |
| WTC | 84 | 0 | 0 | 96 | 0 | 0 | 180 | 0 | 0 |
| **Total** | **651** | **227** | **0** | **652** | **366** | **0** | **1303** | **593** | **0** |
| | **Cases** | | | **Control** | | | **Total** | | |

| Smoking | Non-Smoker | Smoker | Missing | Non-Smoker | Smoker | Missing | Non-Smoker | Smoker | Missing |
|---|---|---|---|---|---|---|---|---|---|
| DNHS | 21 | 18 | 1 | 46 | 14 | 0 | 67 | 32 | 1 |
| GTP | 47 | 27 | 0 | 139 | 52 | 0 | 186 | 79 | 0 |
| MRS | 24 | 39 | 0 | 32 | 31 | 0 | 56 | 70 | 0 |
| PRISMO | 7 | 20 | 5 | 5 | 18 | 7 | 12 | 38 | 12 |
| STARRS | 14 | 37 | 0 | 17 | 34 | 0 | 31 | 71 | 0 |
| TRUST | 67 | 46 | 3 | 158 | 29 | 0 | 225 | 75 | 3 |
| VA-M-AA | 102 | 73 | 8 | 135 | 39 | 12 | 237 | 112 | 20 |
| VA-M-EA | 57 | 30 | 0 | 73 | 16 | 0 | 130 | 46 | 0 |
| VA-NCPTSD | 92 | 42 | 14 | 56 | 7 | 2 | 148 | 49 | 16 |
| WTC | 72 | 12 | 0 | 91 | 5 | 0 | 163 | 17 | 0 |
| **Total** | **503** | **344** | **31** | **752** | **245** | **21** | **1255** | **589** | **52** |

**Please add a footnote that spells out each of the programs so the reader doesn't have to leaf through the text to find what the acronyms stand for.**

**Table S2. Gene Ontology enrichment among top PTSD-associated CpGs (FDR<0.2).**

| | Term | Ont | N | DE | P.DE | FDR |
|---|---|---|---|---|---|---|
| GO:0072678 | T cell migration | BP | 35 | 3 | 1.10E-05 | 2.30E-01 |
| GO:0032010 | phagolysosome | CC | 5 | 2 | 2.22E-05 | 2.31E-01 |
| GO:0005767 | secondary lysosome | CC | 8 | 2 | 6.76E-05 | 4.69E-01 |
| GO:0072676 | lymphocyte migration | BP | 74 | 3 | 1.00E-04 | 5.20E-01 |
| GO:0016191 | synaptic vesicle uncoating | BP | 1 | 1 | 1.31E-03 | 1.00E+00 |
| GO:1901632 | regulation of synaptic vesicle membrane organization | BP | 1 | 1 | 1.31E-03 | 1.00E+00 |
| GO:1903388 | regulation of synaptic vesicle uncoating | BP | 1 | 1 | 1.31E-03 | 1.00E+00 |
| GO:1903390 | positive regulation of synaptic vesicle uncoating | BP | 1 | 1 | 1.31E-03 | 1.00E+00 |
| GO:0071976 | cell gliding | BP | 1 | 1 | 1.51E-03 | 1.00E+00 |
| GO:0072680 | extracellular matrix-dependent thymocyte migration | BP | 1 | 1 | 2.47E-03 | 1.00E+00 |
| GO:0072681 | fibronectin-dependent thymocyte migration | BP | 1 | 1 | 2.47E-03 | 1.00E+00 |
| GO:2000413 | regulation of fibronectin-dependent thymocyte migration | BP | 1 | 1 | 2.47E-03 | 1.00E+00 |
| GO:2000415 | positive regulation of fibronectin-dependent thymocyte migration | BP | 1 | 1 | 2.47E-03 | 1.00E+00 |
| GO:0032127 | dense core granule membrane | CC | 1 | 1 | 2.47E-03 | 1.00E+00 |
| GO:0071133 | alpha9-beta1 integrin-ADAM8 complex | CC | 1 | 1 | 2.47E-03 | 1.00E+00 |
| GO:0000939 | condensed chromosome inner kinetochore | CC | 2 | 1 | 2.57E-03 | 1.00E+00 |
| GO:0006897 | endocytosis | BP | 596 | 5 | 2.60E-03 | 1.00E+00 |
| GO:0071277 | cellular response to calcium ion | BP | 50 | 2 | 2.93E-03 | 1.00E+00 |
| GO:0072679 | thymocyte migration | BP | 2 | 1 | 3.07E-03 | 1.00E+00 |

Gene Ontology terms: "BP" = Biological Process, "CC" = Cellular Component, "MF" = Molecular Function

**Figure S1. Forest Plots of association between PTSD and *AHRR* CpGs in each cohort.**

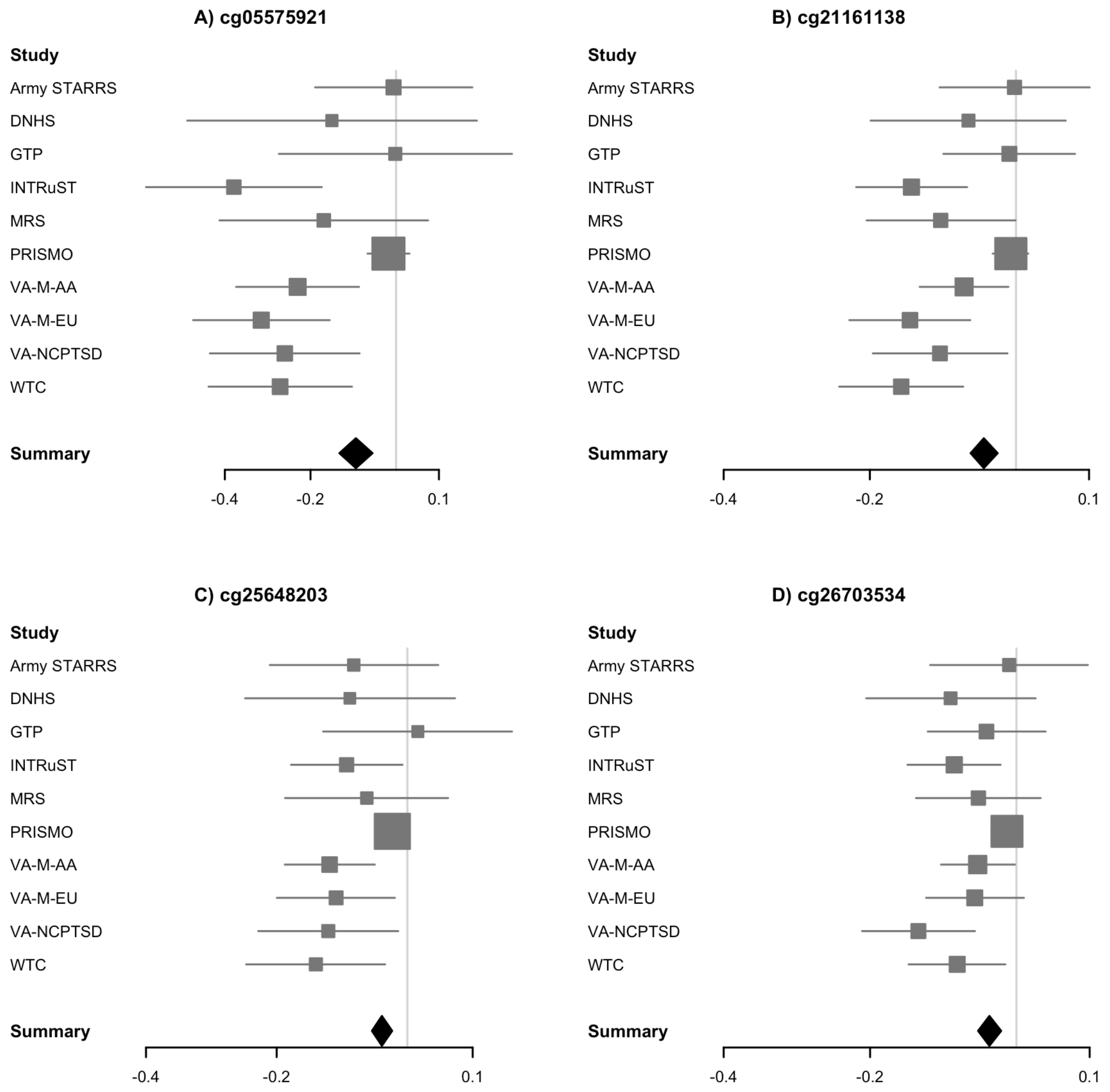

**Figure S2. Comparison of effect sizes before and after controlling for smoking.**

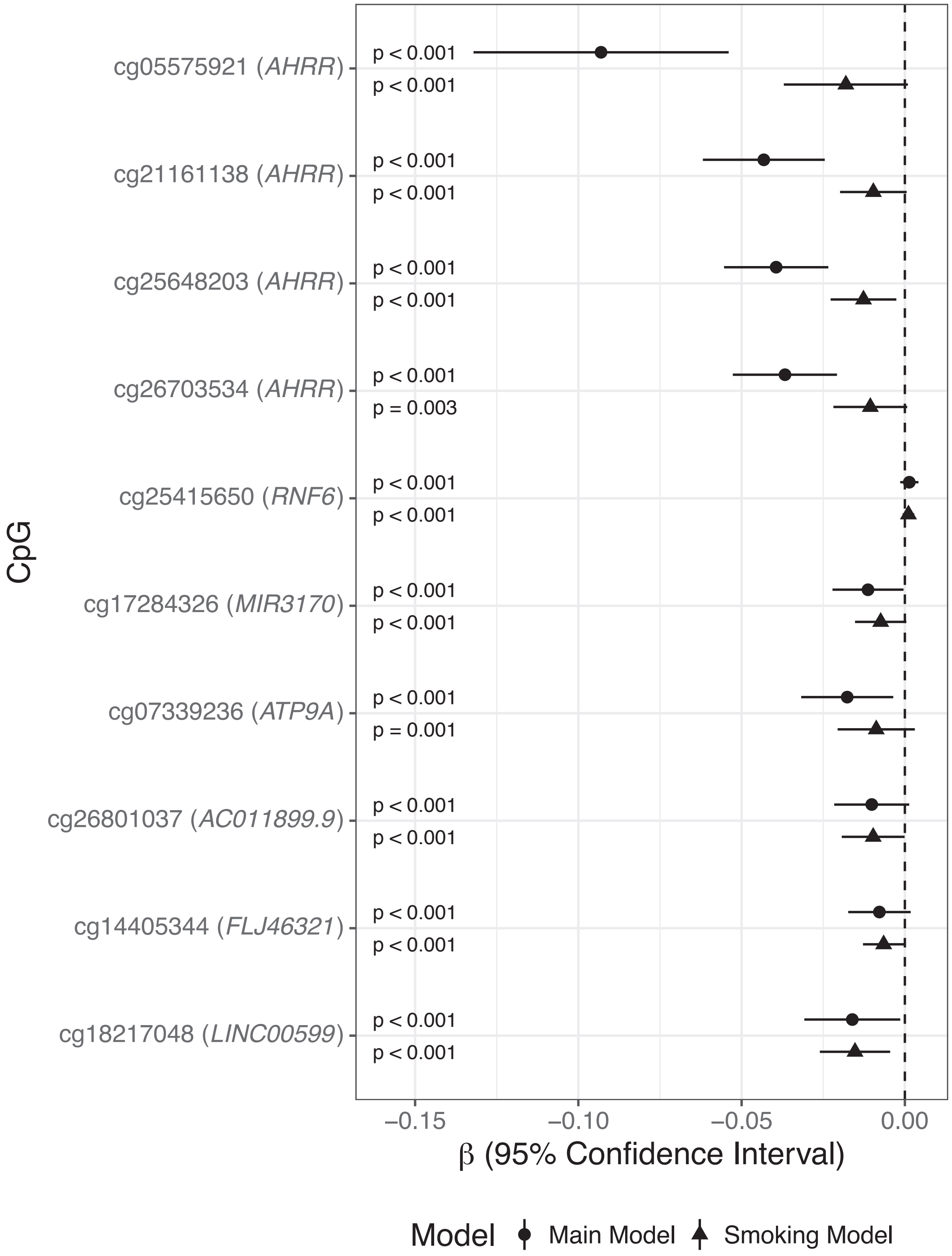

**Figure S3. Association of Smoking-Associated CpG sites with current PTSD in analyses stratified by self-reported smoking status.** The x-axis is the effect size for each association and includes the 95% confidence interval.

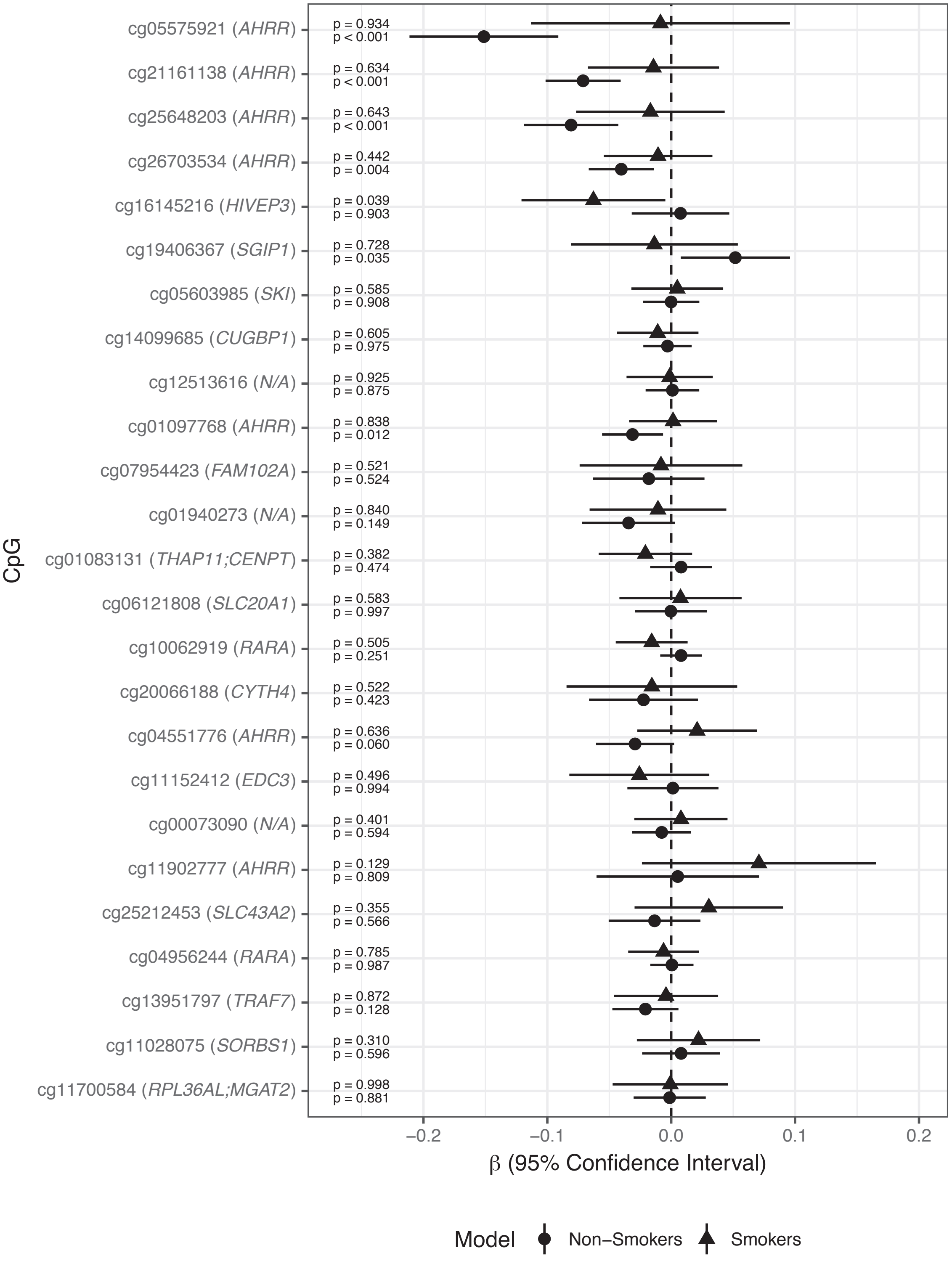

**Figure S4. Lower Kynurenine levels among PTSD cases in the MRS cohort.**

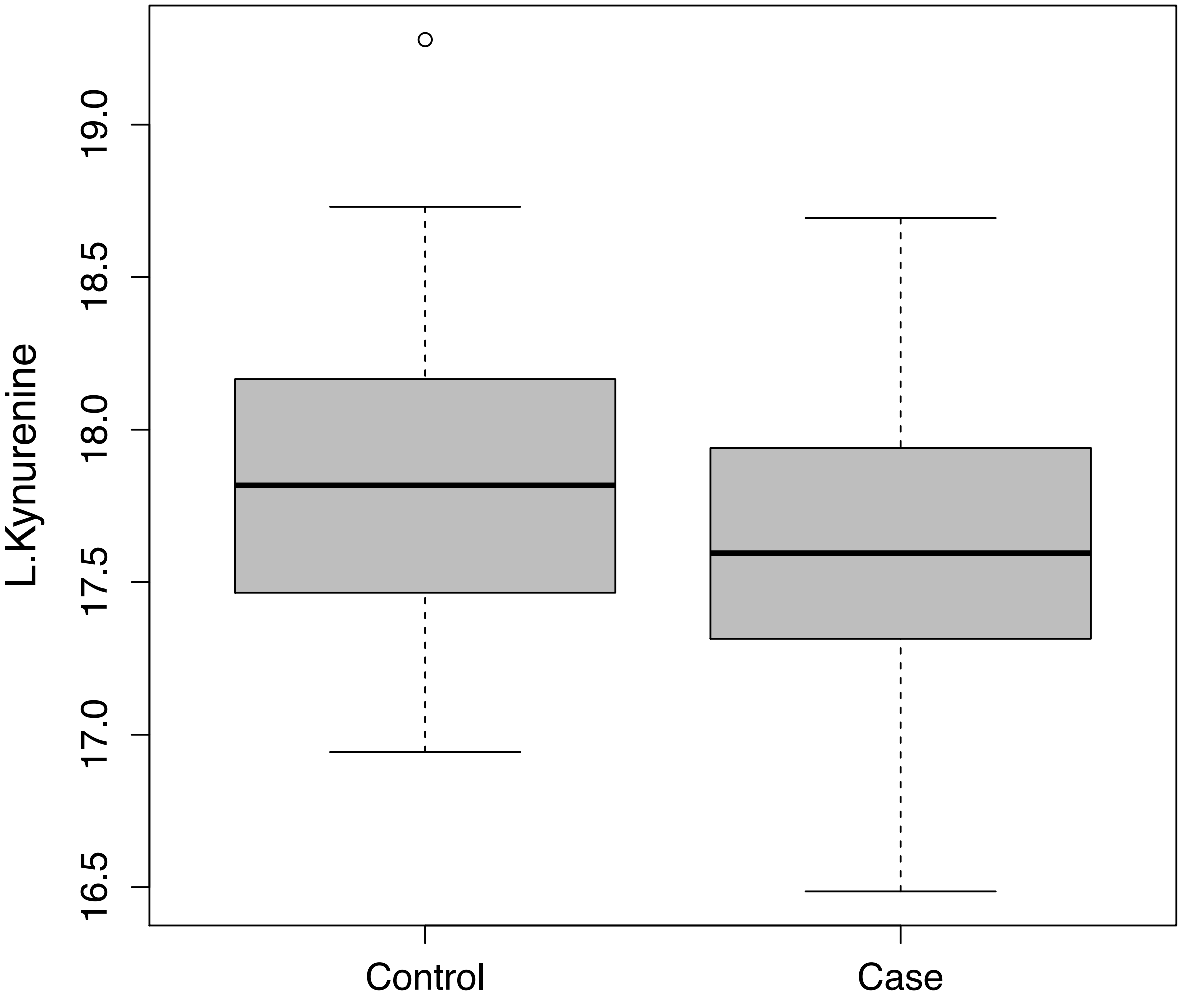

**Figure S5. Correlation of cotinine and kynurenine across all MRS subjects**.

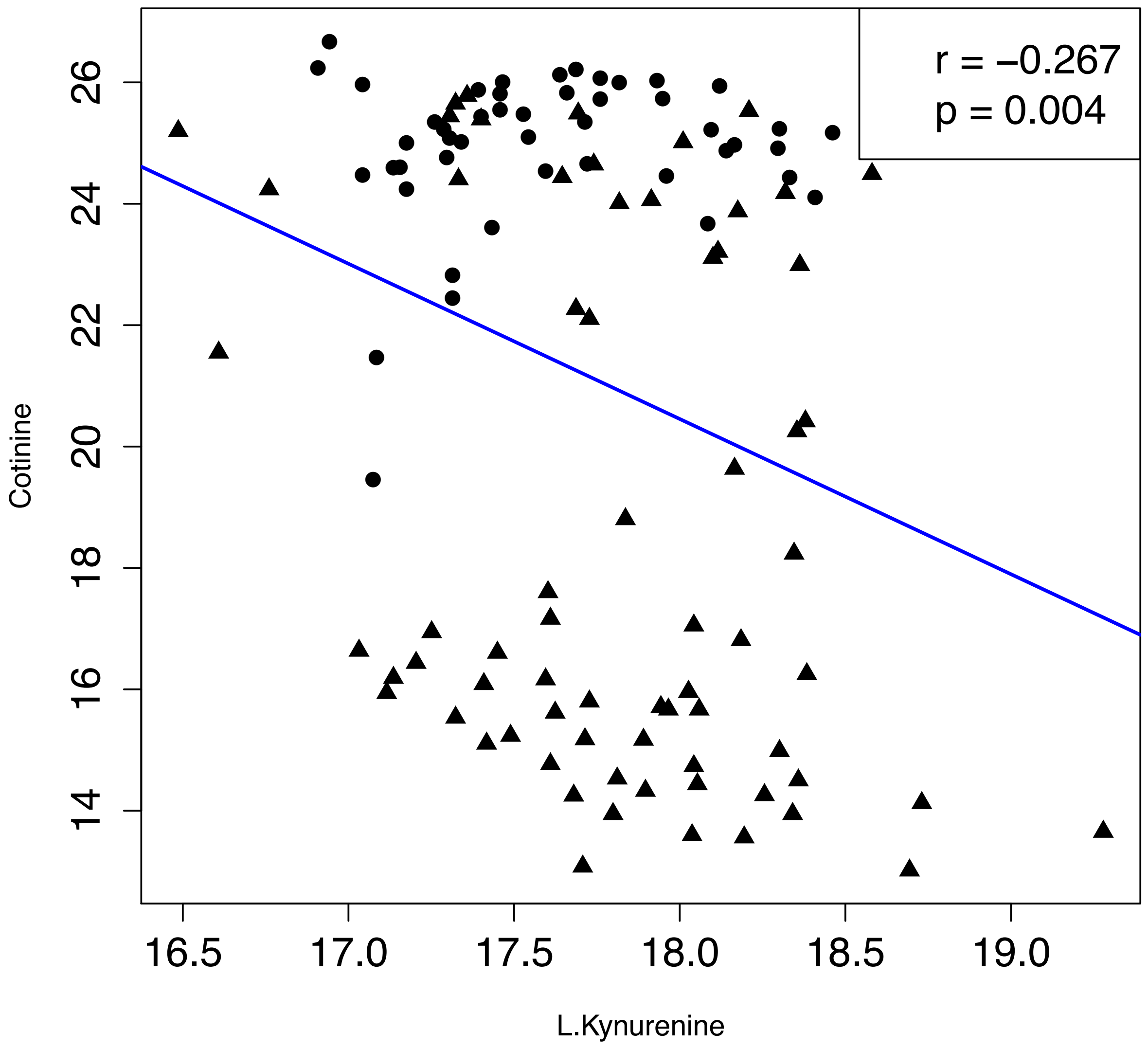
